# Mechanical reinforcement of protein L from *Finegoldia magna* points to a new bind-and-search mechanism

**DOI:** 10.1101/731059

**Authors:** Narayan R. Dahal, Joel Nowitzke, Annie Eis, Ionel Popa

## Abstract

Several significant bacterial pathogens in humans secrete surface proteins that bind antibodies in order to protect themselves from the adaptive immune response and have evolved to operate under the mechanical sheer generated by mucus flow, coughing or urination. Protein L is secreted by *Finegoldia magna* and has several antibody-binding domains. These domains have two antibody-binding sites with vastly different avidity and the function of the second weaker binding interface is currently unknown. Here we use magnetic tweezers and covalent attachment via HaloTag and SpyTag to expose Protein L to unfolding forces in the absence and presence of antibody-ligands. We find that antibody binding increases the mechanical stability of protein L. Using the change in mechanical stability as a binding reporter, we determined that the low-avidity binding site is acting as a mechano-sensor. We propose a novel mechanism where the high-avidity binding site engages the tether, while the low-avidity binding site acts as a mechano-sensor, allowing bacteria to sample the antibody surface concentration and localize its search during successful binding under strain.

**Significance:** It is well known that bacteria have an arsenal of tools to invade and to avoid dislocation. Based on the molecular response of a protein used by anaerobic bacteria to attach to antibodies and disrupt the immune system, here we report on a force-sensor-like behavior, triggered by antibody clusters and force. This pseudo-catch bond between bacteria and antibodies is activated through a second binding site which has lower avidity to antibodies, and which acts as a mechanical sensor, potentially regulating the search radii of the bacterium. Understanding of the bacteria attachment mechanism is of great importance toward developing new antibiotics and mechano-active drugs.

## Introduction

Of critical importance for bacteria to operate in a dynamic environment is to sample the mechanical forces generated by mucus flow, coughing or urination, and adapt accordingly to avoid dislocation or degradation. Bacteria secrete tens to hundreds of multidomain proteins to attach to their host (1). The mechanisms used by bacteria are as diverse as they are fascinating. *(i)* Following the folding of secreted protein domains by *Streptococcus pyogenes* and other Gram-positive bacteria, which contain adjacent carboxyl and amine terminated amino acids, an intramolecular isopeptide bond can cement the folded structure (2). Folded domains with intramolecular isopeptide bonds can withstand nanoNewtons forces, similar to covalent bonds (3), and cannot be unfolded for regular degradation (4). *(ii)* Some *Escherichia coli (E. coli)* secrete α_2_-macroglobulin anti-proteases, which utilizes a Venus flytrap-like mechanism based on the thioester bond, whereby a bait region attracts and inactivates proteases (5). Reformation of the cleaved thioester bond was directly related to force sensing and anchor stabilization (6). *(iii) E. coli* and other bacteria residing in the intestine have also developed a catch-bond based adhesion, where high flow above a threshold generates strong adhesion, while in low or no flow the bacteria can easily detach (7).

An interesting system is that of bacteria secreting antibody binding proteins (ABP) (8). These proteins are binding to plasma or membrane bound antibodies outside the antigen region or secrete ABP. Like a prey turned into predator, binding of ABPs to the most advanced immune molecules is thought to disrupt the immune response and prevent phagocytosis, giving bacteria an evolutionary advantage (9). The exact mechanism and response of these proteins during bacteria adhesion is still poorly understood (10), but thought to be of great importance toward developing new antibiotics and mechano-active drugs (11). From secreted ABPs, protein L stands out as it targets the *κ*-light chain region of antibodies, rather than the heavy chain (12). By targeting the *κ*-light chain region, which is found in ~2/3 of all human antibodies, secreted protein L can bind not only to IgGs, which are responsible with immune memory, but also to IgA, responsible for regulating the microbiota in the mucus, IgM, which is an important part of the initial immune response, or IgE, which could trigger the release of histamine (13). *Finegoldia magna* (formerly *Peptostreptococcus magnus*) secretes protein L as a chain of several domains: a wall domain W, a membrane bound domain M, several C domains (varying depending on the strain), five B domains and one A domain (See also Figure 1A). All B domains have developed binding affinity to antibodies at the *κ*-light chain site and have the residues involved in antibody binding conserved (14). Two binding interfaces were found for antibody-binding of protein L, both targeting the same region of the *κ*-light chain, but with vastly different avidity and unknown function (15).

**Figure 1.**
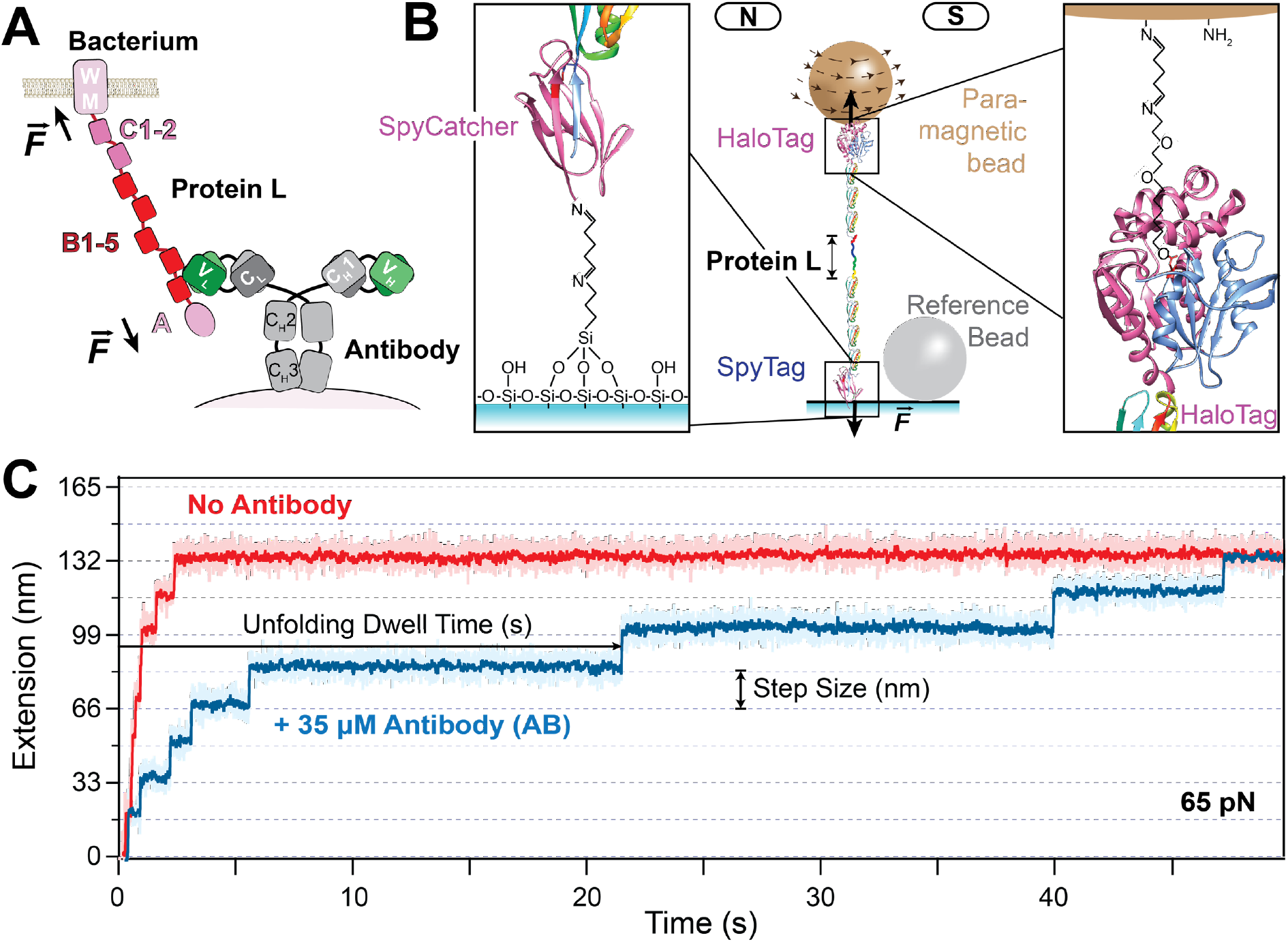
Measurement of antibody binding to protein L using magnetic tweezers and HaloTag-SpyTag covalent attachments. A) Schematics of attachment of a multidomain protein L (top) secreted by *Finegoldia magna* binding to the light chain region of an IgG antibody (bottom) (adapted from ref. (23)). B) Schematics of the tethered polyprotein engineered with a SpyTag and a HaloTag. Inset left: attachment chemistry used for the SpyCatcher/SpyTag reaction to attach the protein to the glass coverslip. Inset right: attachment chemistry used for the chloroalkane-ligand/HaloTag reaction to attach the protein to the amine-terminated paramagnetic bead. C) Example of a trace of the same single molecule unfolding all its eight domains in the absence (red) and presence (blue) of antibodies (35 μM) under a constant force (65 pN). Each step corresponds to the unfolding and extension of a protein L domain. The unfolding dwell-time and step size are defined as indicated by the arrows.

Here we investigate the mechanical response of the B1 domain of protein L (referred to from now on as simply protein L) in the presence of *κ*-light chain IgG antibodies. We find that antibody binding acts as a mechanical sensor for protein L, reinforcing the protein at forces over ~50 pN per molecule. Using this change in mechanical stability as a binding reporter, we measure a binding constant similar to that of the low-avidity binding interface. We propose that this mechanical sensor allows bacteria to sample the local antibody concentration, adjust their search radius and localize their target.

## Results

### A HaloTag-SpyTag based approach for covalent attachment of single proteins

To measure the mechanical unfolding of protein L in the presence of antibody ligands, we use a combination of single molecule magnetic tweezers and covalent attachment. Magnetic tweezers can expose single protein molecules to forces in the pico-Newton range for extensive periods of time, regularly of several hours per molecule (16). Force is applied through the separation between a pair of permanent magnets and a tethered paramagnetic bead and the extension is measured from the displacement of this bead in respect to a reference bead. An unfolding event registers as a nanometer step increase in the end-to-end protein length, with its size dependent on the applied force and number of amino acids inside the folded structure. To achieve these long tethering times, an active focus correction mechanism is used, where a non-magnetic reference bead glued to the glass surface is kept in focus by moving the objective vertically with the help of a piezo actuator. Covalent attachment is desirable, as it results in the most stable tethers and enable longer experiments at higher forces. Several specific covalent chemistries have been developed, based on HaloTag (17, 18), SpyTag (3), cohesin-dockerin (19), and click chemistry (20, 21). For our experiments here, we have engineered eight repeats of protein L sandwiched between a HaloTag at the N-terminus and a SpyTag at the C-terminus (Figure 1B). While previously we used the Biotin-Streptavidin interaction to tether proteins through a C-terminus AviTag (16), this noncovalent attachment becomes challenged when forces above ~60 pN are applied for over 1 minute (22) and could not have been used for the current experiment, where ligand binding increases the mechanical stability of protein L beyond this range. The breaking of the tether at high forces was solved here by using the SpyTag-SpyCatcher link, which can form a covalent isopeptide bond (3). As opposed to the HaloTag-chloroalkane ligand interaction, which forms a covalent ester bond in under 1 second (17), the isopeptide bond formation between SpyTag-SpyCatcher requires several minutes. Hence the glass surface was functionalized with SpyCatcher proteins before it was left to react with our C-terminated SpyTag protein L construct for 30 min (see Methods section for more details). Following a washing step, surface attached proteins were left to react for ~1 minute with the chloroalkane terminated superparamagnetic beads at the HaloTag site, before the magnets were brought down. This time is more than sufficient for the HaloTag interaction, and avoids non-specific or multiple tethers between the bead and the surface, which could form if longer times would be allowed for this step. When a force of 65 pN is applied to our protein L construct, we measure eight equidistant unfolding steps, unraveling in ~3 s (Figure 1C). The HaloTag-SpyTag attachment also allows us to change the solution buffer inside the fluid chamber without breaking the molecular tether, and enabling the measurement of the same protein molecule in different concentrations of antibody. When we exchange the antibody-free solution buffer with one that contains *κ*-light chain antibodies and apply the same 65 pN to the same molecule, we find that ~47 s are now needed to completely unfold all protein L domains (Figure 1C, blue trace). Hence, the antibody binding has a mechanical strengthening effect on protein L, and this effect can also be used to measure the binding of antibodies.

### Using mechanical unfolding to measure binding of protein L to antibodies

To measure the binding interaction between protein L and IgG antibodies, we use a two-step force pulse protocol, which allows us to determine how many domains have a ligand attached (Figure 2). First, the force is ramped to a low-force (45 pN) and maintained at this value for a total of 35 s. At this force, the unfolding rate of protein L is 0.25 ± 0.01 s^−1^ (it takes on average ~4 s dwell time to unfold a domain) and the exposure time is generally sufficient to unfold all the protein L domains free of antibodies. We then ramp the force once more and maintain it at 100 pN for 100 s. This second high-force pulse is used to determine the number of protein L domains with bound antibodies. Indeed, without any antibodies added, protein L unfolds all its eight domains in the first low-force pulse (45 pN, Figure 2A). When the protein L is measured in a solution containing 35 μM antibodies, most of the unfolding events appear in the high-force pulse (100 pN). At the end of each 100 s – 100 pN exposure, the protein is left to refold at ~2 pN for 100 s and bind new antibody molecules from solution. As we can tether single protein L molecules for extensive times and expose them to alternating high and low force pulses, we are effectively resetting the binding process with every cycle. We then quantify the binding as the number of unfolding domains in the 100 pN region over the total number of domains (the last bar in Figure 2C). In 35 μM IgG, ~75% of the unfolding events appear in the high-force 100 pN pulse (Figure 2C).

**Figure 2.**
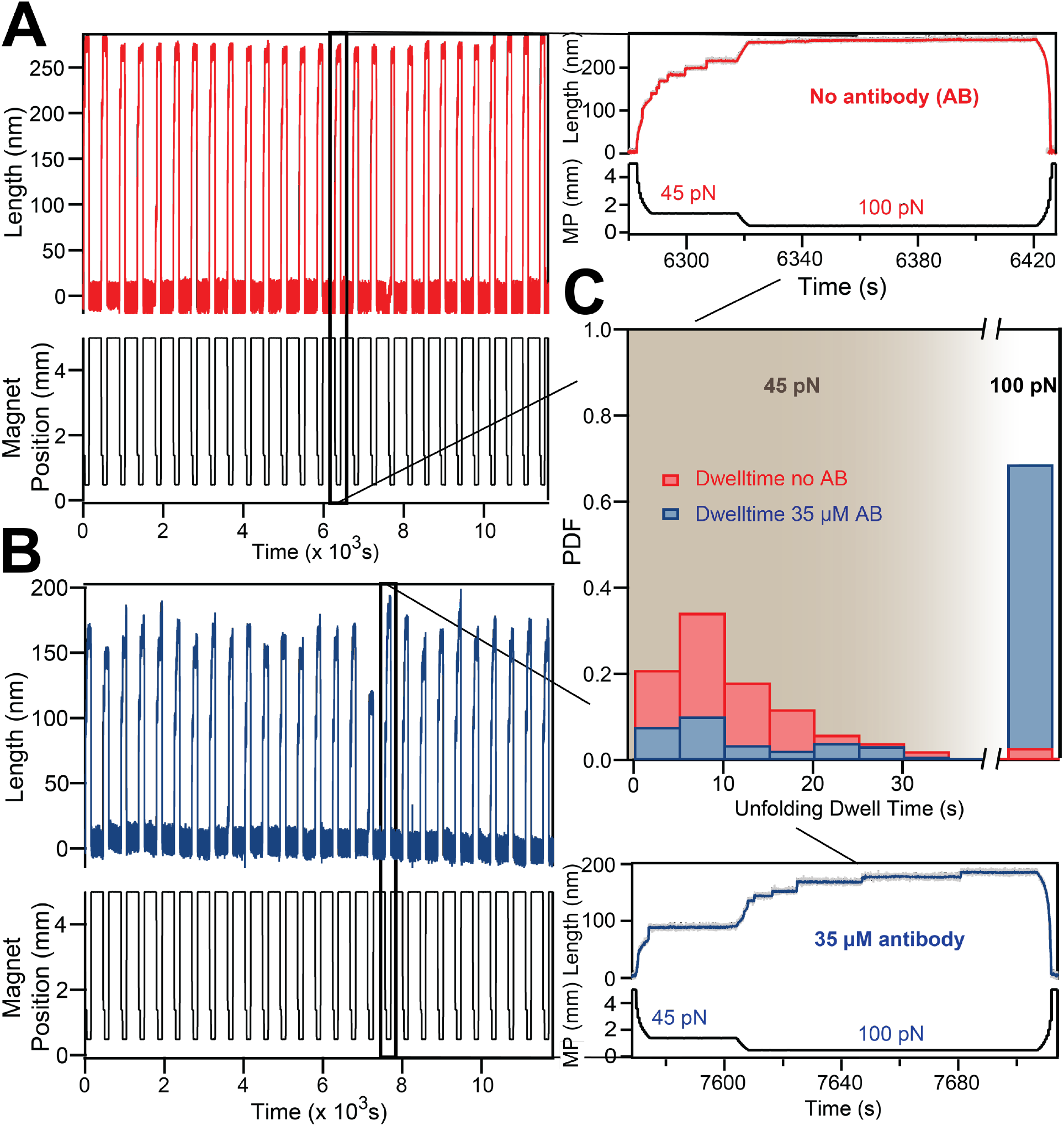
Measurement of antibody binding using a two-step protocol. A) Representative unfolding trace of octamer of protein L domain in absence of antibody. The force protocol was set to 45 pN for ~35 seconds, followed by ramping the force to 100 pN. Zoom in (Top right) shows the unfolding of all 8 domains within 30 seconds at 45 pN. B) Similar unfolding trace obtained from the same construct with same force protocol measured in the presence of 35 μM mouse serum IgG. Zoom in (Bottom right) shows the unfolding of the majority of protein domains at high force (100 pN) in the presence of antibody. C) Unfolding dwell time frequency histograms of protein L domains in absence and in presence of 35 μM IgG. In absence of IgG, more than 90% of domains unfold within 35 seconds at the low pulling force of 45 pN (red histogram) whereas in the presence of 35 μM IgG, most of the domains unfold in the high force pulse of 100 pN (blue histogram).

By repeating the two-pulse protocol with changing antibody concentrations, we can determine the binding constant to protein L (Figure 3A). As the antibody concentration is increased, more and more unfolding events appear in the 100 pN part of the pulse. However, the binding probability plateaus at a value of ~0.75 at concentrations above 30 μM (Figure 3B). The fitted dissociation constant between the IgG antibodies and protein L, using the Hill-Langmuir equation *X*_*AB-L*_ = [*AB*]/(*K*_*D*_ + [*AB*]), has a value of 23 ± 3 μM.

**Figure 3.**
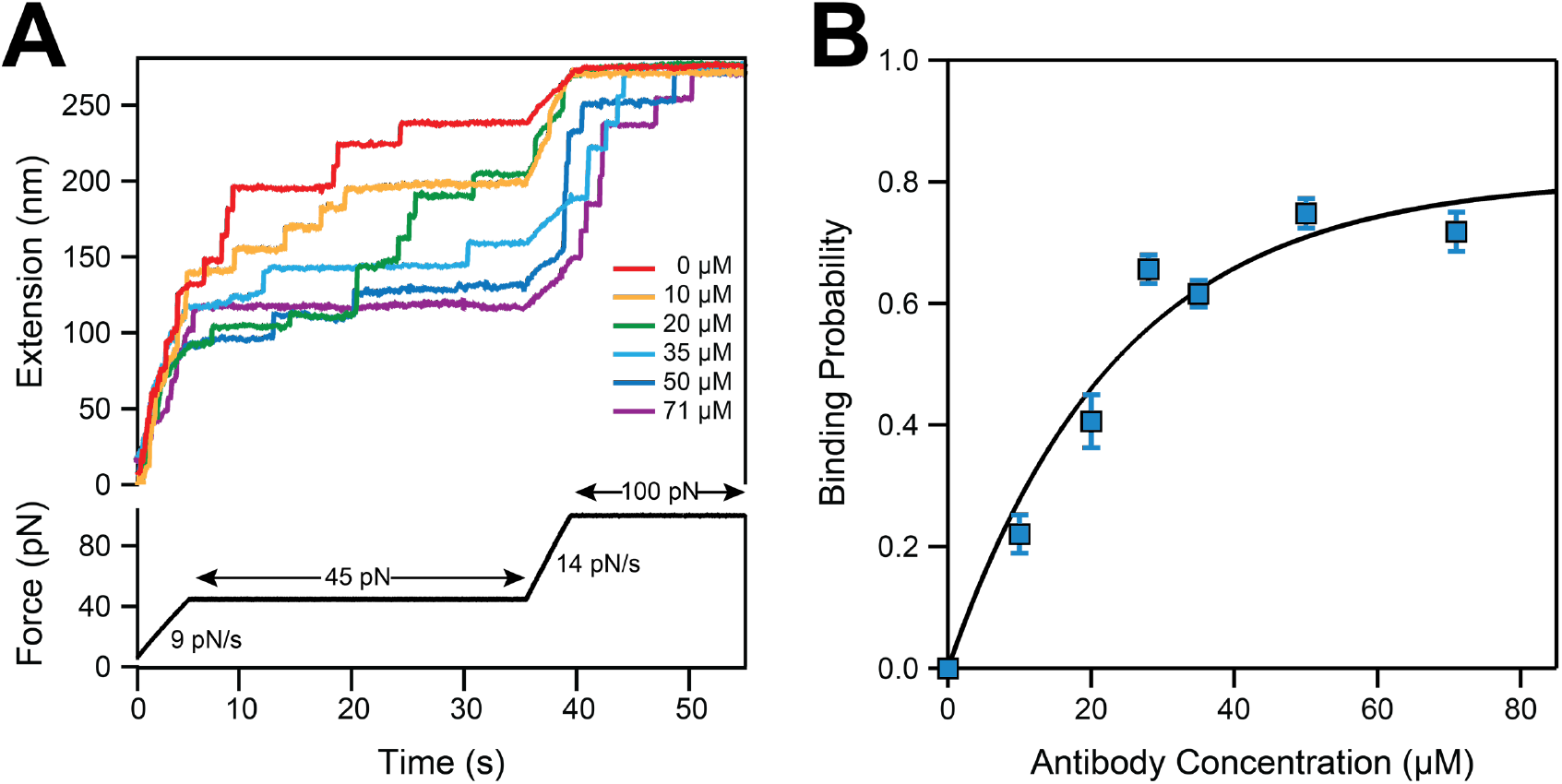
Determining the dissociation constant from the change in the mechanical stability of protein L. A) Unfolding traces of protein L octamer from the HaloTag-(protein L)_8_-SpyTag construct, measured in different concentration of mouse IgG antibody. Without antibody, all the domains unfold at low force (45pN, red trace) whereas a high concentration of antibody requires a high force (100 pN, violet trace, 71 μM antibody) to unfold. B) The binding probability as a function of the concentration of IgG. Increasing the concentration of antibody increases the binding probability and thus the stability. Blue squares represent the binding probability at different concentration of antibody. The line represents a fit using Hill-Langmuir equation and yielding a dissociation constant *K*_*D*_ = 23 ± 3 μM.

### Antibody-binding induces a pseudo catch-bond behavior in protein L unfolding under force

While our single molecule assay constitutes an elegant approach to measure antibody binding, it also can determine the unfolding kinetics of protein L in the presence and absence of its IgG ligand. For measuring unfolding kinetics, we use the square-root histogram method (24, 25) (histogram of logarithmic binning of the unfolding dwell time – see Figure 4A and B). In this case, the protein L octamer construct was exposed to a single constant force in the absence and presence of antibodies at a saturating concentration (see also Figure 3B). The dwell time for unfolding was determined as a function of force and experimental conditions, as defined in Figure 1C. Histograms were then constructed from the natural logarithm of the measured dwell-times and fitted to a single-exponential law: exp [*x* − *x*_0_ − *exp*(*x* − *x*_0_)] when a single peak was present, and a double exponential law: A_1_exp [*x*_1_ − *x*_01_ − *exp*(*x*_1_ − *x*_01_)] + A_2_exp[*x*_2_ − *x*_02_ − *exp*(*x*_2_ − *x*_02_)] when the histogram had two peaks (with *x* = ln [*t*], *x*_0_ = − ln[*r*(*F*)] where *t* is the unfolding dwell time and *r*(*F*) is the force-dependent unfolding rate. The square-root histogram method has the advantage of separating processes taking place on different characteristic timescales. The distribution of unfolding events at low forces exhibited a bimodal shape with ~ 10-20 % of the events in a weak state (black points in Figure 4C) and the remaining in a more mechanically stable state. This behavior was attributed to ephemeral states and domain swapping in a previous study (25). As the experienced force is increased, the histogram peak of the unfolding dwell times moves to lower dwell-time values and the first peak is no longer present (compare red histograms in Figures 4A and B).

**Figure 4.**
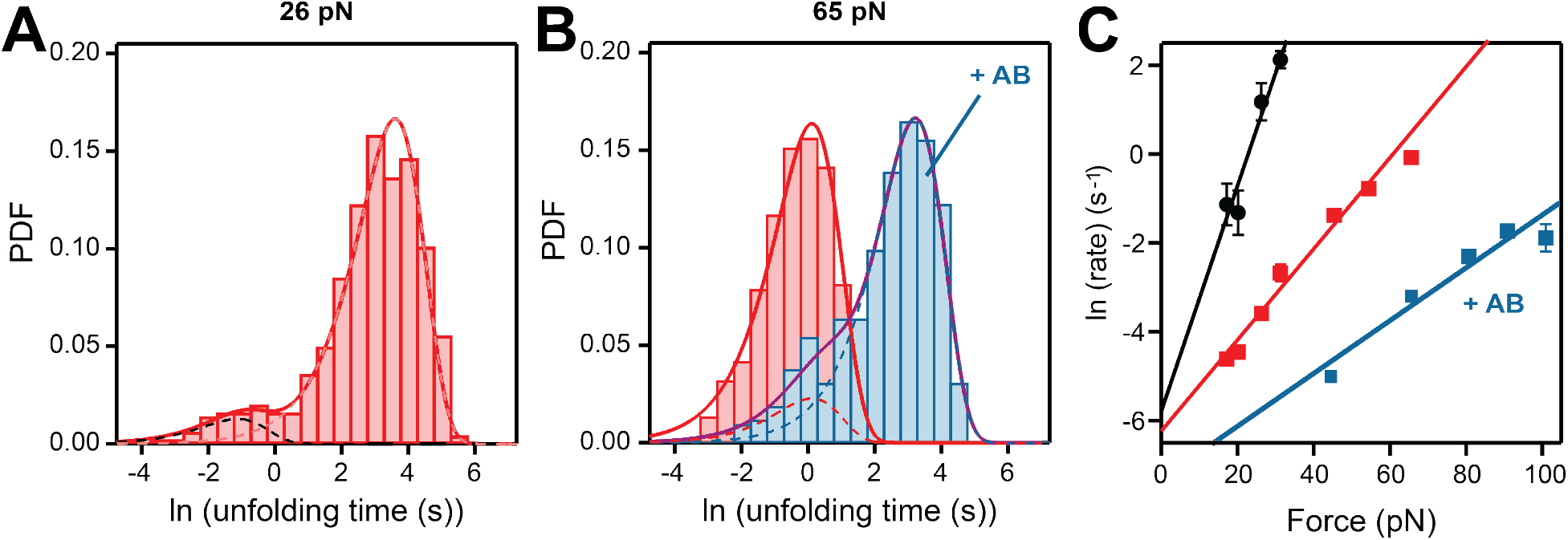
Force-dependent unfolding kinetics of protein L in the presence and absence of antibodies. A) Histogram of the natural logarithm of the measured dwell-times of protein L without added antibodies, at 26 pN. The dotted lines represent the individual fits using a single exponential law, while the continuous line is their sum. Between 10-20% of protein L domains are measured in a mechanically weak state, a number similar to the percentage of domains that do not bind antibodies, but a direct correlation between the two populations cannot be readily made. This weak state was previously attributed to domain swapping (24, 28). B) Histogram of the natural logarithm of the measured dwell-times of protein L at 65 pN without antibodies (red) and in the presence of antibodies (35 μM, blue). The first peak in the blue histogram coincides with the location of the red peak and has an amplitude that corresponds to ~12% unbound domains, in agreement with the experiments from the double-pulse protocol (Figure 3). C) Unfolding rates of the weak state of protein L (black circles), of the native state (red squares), and antibody-bound state (blue squares). The lines represent the fits using the Bell’s model.

The square-root histogram method is very useful at high forces (>40 pN) for separating the unfolding events of protein L arising from domains that have bound antibodies from the ones that do not (Figure 4B). In this case, we first use the antibody-free experiments to determine the unfolding rates at a given force (red histogram Figure 4B). We then fit a double exponential law to measure the unfolding kinetics of antibody-bound protein L domains (blue histogram Figure 4B). To describe the unfolding rates as a function of force, we then use the Bell model 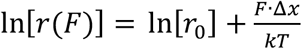 (Figure 4C), where *r*_0_ is the extrapolated rate at zero force, *F* is the applied force, Δ*x* is the distance to transition state and *kT* the Boltzmann thermal energy. An interesting finding is that the unfolding rate of the protein L domains with bound antibodies has a different dependency slope with force than the unfolding rate of the protein L free of antibody (blue vs red points in Figure 4C). These dependencies are characterized by a distance to transition state of 0.24 ± 0.04 nm for protein L with bound antibody and 0.42 ± 0.01 nm for protein L without bound antibodies, and suggests that the higher the experienced force, the larger the mechanical stabilization effect is.

## Discussion

Protein L is secreted by *Finegoldia magna* to attach to antibodies, and has an α-β conformation (from N-to-C: β1-β2-α-β3-β4). The B1 domain of protein L has between 61 to 89% sequence homology with the B2-5 domains and the binding interfaces are highly conserved (14). *Finegoldia magna* was found in vastly diverse biomechanical environments, on mucous membranes of the mouth, upper respiratory, gastrointestinal, and genitourinary tracts (26), and must withstand mechanical stress to prevent being removed by fluid flow (27) (Figure 5A). Two key findings result from our single molecule measurements, which will be discussed below: (i) the low avidity binding site is responsible with mechano-sensing, while the high-avidity site does not influence the mechanical stability of protein L; (ii) the mechanical activation of protein L is reminiscent of a catch-bond, where the larger the experienced force, the bigger the difference in stability between protein L with bound antibodies versus protein L alone.

**Figure 5.**
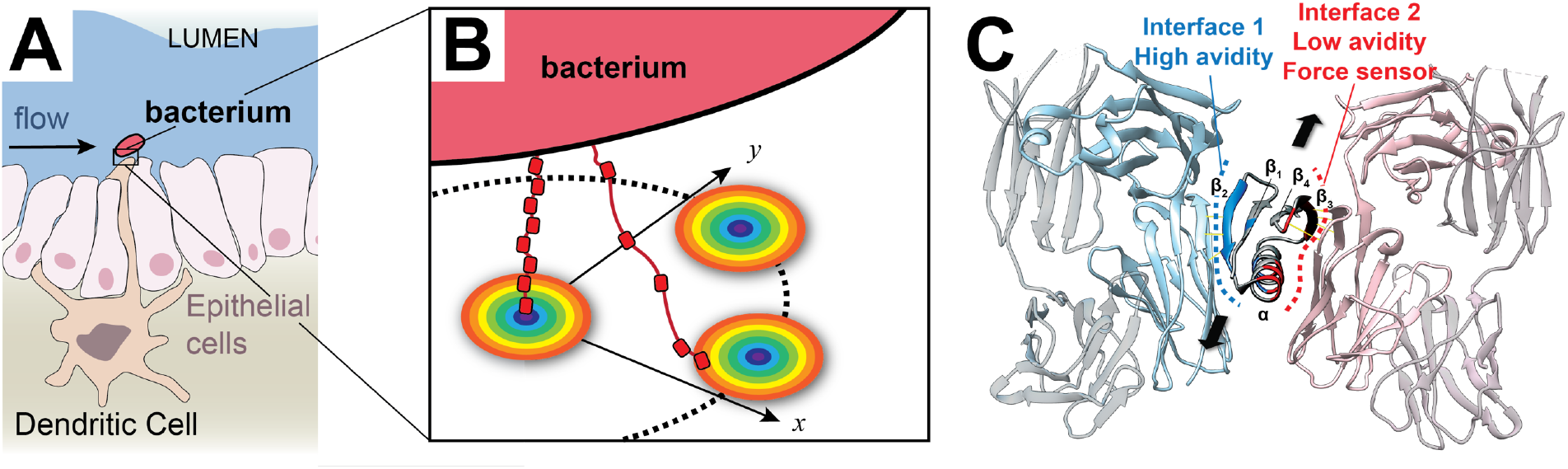
Proposed force-activated mechanism for bacteria adhesion. A) Schematics of a dendritic cell presenting a IgG covered surface inside the lumen, where opportunistic pathogen such as *Finegoldia magna* (red) can attach under a mucus flow. B) Proposed mechanism, where the bacterium secretes protein L multidomains to attach to antibodies. The circles denote the antibody clusters present at the cell surface (29). High antibody concentrations will lock protein L in a folded conformation by populating both interfaces, reducing the search radius. Low antibody concentration will allow attachment at the high-avidity interface, without affecting the mechanical stability and increasing the search radius. C) Ribbon representation of protein L bound to two antibody molecules. The high-avidity interface is shown in blue, while the low-avidity interface, which can act as a force-sensor, is drawn with red (based on PDB: 1HEZ (15)). The arrows show the direction of the force vector.

Using the change in mechanical stability of protein L, we measured a binding constant to antibodies of 23 ± 3 μM. The measured binding constant is smaller than that reported from titration experiments, which was 0.1-0.2 μM (28). The same authors reported that treatment with tetranitromethane, which is a tyrosine inhibitor, prevents normal antibody biding at the β1-β2-α interface, and decreases the binding constant to ~30 μM (28). This change in binding affinity was later explained by the discovery of a second binding interface at the α-β3 site (15). The measured value here for the binding constant via mechanical unfolding suggests that it is this second interface that plays a role in mechanosensing.

Taken together, our results point to a novel pseudo catch-bond mechanism. It is well-known that antibodies form transient clusters on the membrane of dendritic cells, when acting as docking sites for the complement system or phagocytes (29). When the bacterium attaches to its substrate, if the antibody surface concentration is low, the high-avidity binding site is more likely to engage, without influencing the mechanical stability of protein L. In this case, the anchored bacteria can unfold and extend its domains and increase its search radius (Figure 5B). When interacting with an antibody cluster, some protein L domains would bridge two antibody molecules at their light-chain region, increasing their mechanical stability and acting as force-sensors (Figure 5B &C). Under flow, when the bacterium engages an antibody cluster, its search radius reduces from ~19 nm/domain to ~4 nm/domain. This reduction in the search radius would allow the bacterium to counteract an immune response. So while the first binding site acts as an attachment ligand due to its high avidity, it is the second binding site that can engage under flow and produce a mechanical signal, informing on the concentration of the antibodies at the target site. We propose that this mechano-sensor will have a significant role in tuning the search distance of bacteria under force and to orient the secreted protein L chains toward either a fight or flight mechanism.

This double site mechanism might also be common in other pathogens. Antibody-binding protein G secreted by group G *streptococci*, has a similar β1-β2-α-β3-β4 structure and attaches IgGs at the heavy-chain region. When measured under force, the binding constant of protein G at the Fc antibody region was also found to be much smaller than that compared from bulk experiments (30), but a second binding interface is not currently known for this complex. Similarly, *Staphylococcus aureus*, which secretes antibody-binding protein A and the clumping factor A was shown to form aggregates under high shear conditions (31). For the clumping factor A, two distinct binding sites were identified, with their adhesion tightly regulated by mechanical force (32). Taken together with our findings here, the double-binding site mechanism might be an important feature used by bacteria to both attach to their target and sample the transient forces, allowing it to better adapt and migrate. Reminiscent of attachment operating under a catch-bond mechanism, this flow-induced search can allow bacteria to selectively engage ligand clusters.

## Methods

Unless otherwise specified, all chemicals were purchased from Sigma-Aldrich. Eight-repeats of protein L were inserted into a modified pFN18a vector (Promega), which introduces a HaloTag at the N-terminus and a SpyTag at the C-terminus. Proteins diluted to ~ 100 nM were left to adsorb on a functionalized SpyCatcher surface for ~30 min, to allow for the maturation of the attachment through an isopeptide bond (3). After washing the non-adsorbed proteins, paramagnetic beads (Thermo Fisher Scientific) with imbedded chloroalkane ligands (Promega) were left to react with the HaloTag end, which results in the formation of a covalent ester bond. Further details on protein engineering, expression and purification, and on surface and bead functionalization are provided in the SI.

The extension of single protein molecules at varying forces were obtained using the magnetic tweezers technique described in refs. (16, 33). Briefly, the chamber was mounted on top of an inverted microscope (Olympus IX71), and the separation between the paramagnetic beads and a pair of permanent magnets was achieved using a voice-coil actuator (Equipment Solution). ROIs of 128×128 pixels were selected around a tethered super paramagnetic bead and a glued non-magnetic reference bead. At the beginning of each experiment, a stack library was obtained for the two selected beads by changing the focusing position with the help of a piezo actuator (P-725, PI) in equal steps of 20 nm. Two-dimensional fast Fourier transforms (2D-FFTs) of the ROI images were then used to obtain a radial profile as a function of focal distance for the two beads. During an experiment, the correlation between the radial profile of each bead was computed against its stack library and a Gaussian fit was used around the maximum of the correlation curve to determine the location of each bead. The extension of the molecule was measured as the difference between the position of the paramagnetic and reference beads. During measurements, any instrumental drift was also corrected by adjusting the position of the objective using the piezo actuator, such that the reference bead was maintained at the same focal point. All data acquisition and processing was done in Igor Pro (WaveMetrics). Data analysis and errors estimation were done as explained in the SI section.

## Acknowledgements

We acknowledge all members of the Popa lab for useful discussions. This work was supported by an NSF Career award (grant number MCB-1846143) and a Shaw award (from Greater Milwaukee Foundation).

## Author contributions

I.P. designed research; N.D. and J.N. performed research; A.E. contributed to new reagents; N.D., J.N. and I.P. analyzed data; and I.P. wrote the paper with input from all authors.

The authors declare no conflict of interest.

